# Photoprotective melanin is maintained within keratinocytes in a storage lysosome

**DOI:** 10.1101/2024.02.05.578910

**Authors:** Matilde V. Neto, Michael J. Hall, João Charneca, Cristina Escrevente, Miguel C. Seabra, Duarte C. Barral

**Author notes:** These authors contributed equally to this work (Matilde V. Neto, Michael J. Hall). Correspondence: Duarte C. Barral, NOVA Medical School, Faculdade de Ciências Médicas, Universidade NOVA de Lisboa, Campo dos Mártires da Pátria, 130; 1169-056 Lisboa, Portugal.

## Abstract

In the skin, melanin is synthesized by melanocytes within melanosomes, and transferred to keratinocytes. After being phagocytosed by keratinocytes, melanin polarizes to supranuclear caps that protect against the genotoxic effects of ultraviolet radiation. We provide evidence that melanin-containing phagosomes undergo a canonical maturation process, with the sequential acquisition of early and late endosomal markers. Subsequently, these phagosomes fuse with active lysosomes, leading to the formation of a melanin-containing phagolysosome that we named melanokerasome. Melanokerasomes achieve juxtanuclear positioning via lysosomal trafficking regulators Rab7 and RILP. Mature melanokerasomes exhibit lysosomal markers, elude connections with the endo/phagocytic pathway, are weakly degradative, retain undigested cargo and are likely tethered to the nuclear membrane. We propose that they represent a lysosomal-derived storage compartment that has exited the lysosome cycle, akin to the formation of lipofuscin in aged cells and dysfunctional lysosomes in lysosomal storage and age-related diseases. This storage lysosome allows melanin to persist for long periods, where it can exert its photoprotective effect efficiently.

## Introduction

Skin pigmentation is a function of epidermal-melanin units, which are composed of approximately one melanocyte per 30-40 keratinocytes (Cichorek et al., 2013). Melanocytes are the producers of the photopigment melanin. Melanin is synthesized and packaged into lysosome-related organelles (LROs) termed melanosomes, but it exerts the bulk of its photoprotective effect in keratinocytes. Therefore, melanin produced in melanocytes must be transferred to keratinocytes and efficiently processed therein. Although several transfer mechanisms have been proposed in the literature, there is now solid evidence that coupled exo/phagocytosis of the melanosome core (*i.e.*, melanocore) is the predominant mode of melanin transfer in basal conditions, in the human skin epidermis (Tarafder et al., 2014). In this process, melanosomes fuse with the melanocyte plasma membrane in a Rab11b- and exocyst-dependent manner, and membrane-less melanocores are subsequently phagocytosed in a Rac1- and Cdc42-dependent manner (Tarafder et al., 2014; Moreiras et al, 2020). After phagocytosis of melanocores by keratinocytes (Moreiras et al., 2022), melanin-containing compartments are trafficked to the juxtanuclear region and form supranuclear caps that function akin to a “parasol”, protecting the nuclear DNA from harmful ultraviolet (UV) radiation. There are variations in the phagocytic process depending on skin color. In dark skins, large individual melanin granules are found within keratinocytes. In contrast, in light skins, smaller granules of melanin organized in clusters are present in keratinocytes, suggesting that the skin phototype is influenced by the differential processing of melanin (Minwalla et al., 2001).

Despite the clear importance of melanin processing in determining and maintaining skin pigmentation, the cell biology and mechanisms underlying this process remain enigmatic. Recently, we demonstrated that melanocore internalization by keratinocytes occurs by phagocytosis, leading to the formation of a melanin-containing storage compartment, which we proposed to be named melanokerasome (MKS) (Moreiras et al., 2021;. Bento-Lopes et al., 2023). MKSs were found by us and others to be weakly acidic and weakly degradative (Correia et al., 2018; Hurbain et al., 2018). We therefore proposed that they could represent either a hybrid compartment or transitional early-to-late endosomal organelles, whose maturation along the endolysomal pathway had been arrested (*i.e.*, evading lysosomal fusion) (Correia et al., 2018). With recent advances in the field, it is now appreciated that lysosomes can cycle through different stages in regard to pH and acid hydrolase activity, and thus can exist in a weakly acidic and poorly degradative state (Bright et al. 2016; Barral et al., 2022). Hence, we postulated that MKSs are a lysosomal compartment with special characteristics that are both unique and vital to skin pigmentation.

Herein, we characterized the melanin-containing compartment in keratinocytes using an experimentally tractable *in vitro* model. Our data suggest that MKSs are neither hybrid or maturation-arrested compartments, but rather mature by acquiring late endolysosomal markers and fusing with active lysosomes. Furthermore, our data suggests that MKSs utilize the lysosomal trafficking machinery for perinuclear positioning to assist the formation of supranuclear caps. Thus, we uncovered a physiological example of lysosomes used as a storage compartment.

## Results

### Melanin-containing compartments associate with early and late endosomes/lysosomes

Initial characterization of MKSs was performed at a single (fixed) time point and, thus, does not provide information about the dynamics of the maturation process and the sequence of events. To shed light on these, we designed an assay to determine whether the early and late endosomal markers are sequentially acquired over time after internalization, which would be consistent with trafficking along the endolysosomal pathway. For this, XB2 keratinocytes were initially incubated with melanocores on ice, to allow the close contact of melanin with the cell surface, but with no/minimal internalization. Phagocytosis of melanocores was subsequently allowed by placing the cells at 37°C. This assay will subsequently be referred herein as a “synchronized pulse” of melanocores. We selected timepoints of 1h, 4h and 24h after internalization to assess the transition from early to late endocytic markers at short, medium and long time points, respectively. After phagocytosis, we observed a small but gradual decrease (∼30% at 24h) of the early endosomal marker EEA-1 puncta surrounding phagocytosed melanocores (Figure 1A and B). However, with time, there is a gradual and significant increase of the co-localization of phagocytosed melanocores with the late endosomal and lysosomal markers, CD63 and LAMP1 (increases of up to 26% and 38%, respectively) (Figure 1C, D and E). These observations are consistent with the hypothesis that melanin-containing phagosomes undergo maturation along the endolysosomal pathway inside keratinocytes.

**Figure 1:**
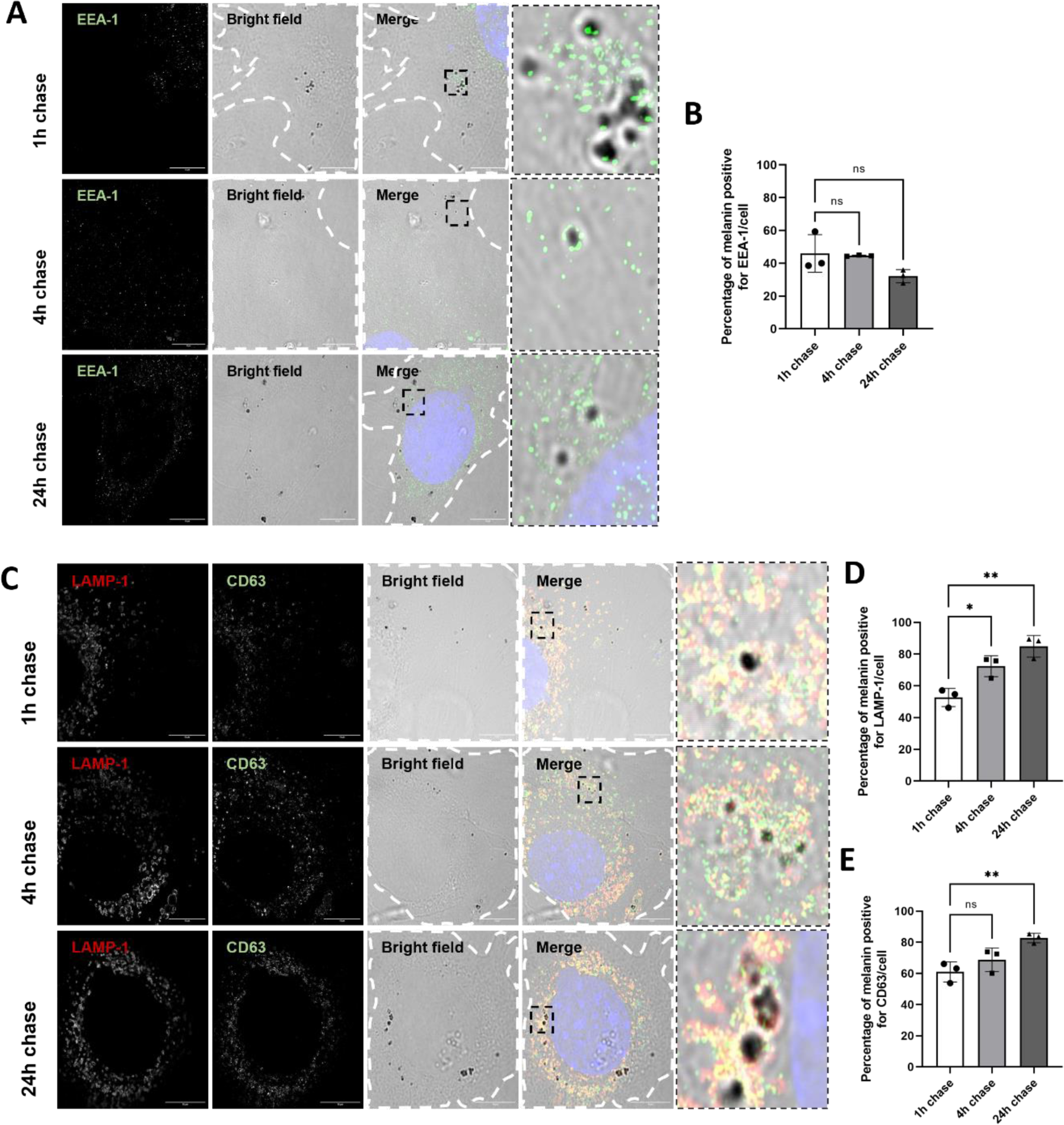
Association of melanin-containing compartments with LAMP1 and CD63 increases over time after phagocytosis. XB2 keratinocytes were incubated with melanocores for 1 hour, washed, and melanin chased for 1, 4 or 24 hours. After the chase, cells were fixed and early endosomes identified by the presence of EEA-1 **(A and B),** and late endosomes/lysosomes by CD63 and LAMP-1 labeling **(C, D and E)**, using immunofluorescence. Nuclei were visualized by DAPI staining (blue). Scale bars = 10 μm. p values (one-way ANOVA) were considered statistically significant (*) and (**) when < 0.01 and < 0.05, respectively, or non-significant (ns) when ≥0.05. Plots show mean ± SD of three independent experiments.

### Melanin-containing phagosomes fuse with pre-existing lysosomes

We previously proposed to name the mature melanin-containing compartment in keratinocytes melanokerasome (MKS) (Moreiras et al., 2021;. Bento-Lopes et al., 2023). After establishing that this compartment is positive for late endosomal and lysosomal markers using relatively long chase periods (up to 24 hours), we next aimed to address whether MKSs are formed by fusion with a pre-existing lysosomal compartment. Using transmission electron microscopy (TEM), we first incubated XB2 keratinocytes with a bovine serum albumin (BSA)-gold conjugate to load the lysosomal compartment, before feeding with melanocores for 2h. Within the 2h pulse, organelles with the lamellar morphology typically associated with lysosomes that also contained aggregated gold could be observed fusing with melanin-containing vesicles (Figure 2A). We then moved to Reconstructed Human Pigmented Epidermis (RHPE) as an in vitro human model system, which we previously showed to mimic the essential features of human pigmentation (Hall and Lopes-Ventura et al., 2022). We observed by TEM similar examples of melanin within compartments with lysosomal morphology (Figure 2B). We further analyzed the MKS compartment in keratinocytes within RHPE by serial section transmission electron tomography to more closely visualize MKS content. We identified melanin within juxtanuclear single-membraned MKSs, also containing other membranes that likely represent lysosomal cargo such as intralumenal vesicles (ILVs) and lysosomal lamellae (Figure 2C). Tomography also revealed a striking close apposition between the outer nuclear envelope membrane and the MKS limiting membrane, which could be maintained by tethers (Supplementary Figure S2).

**Figure 2:**
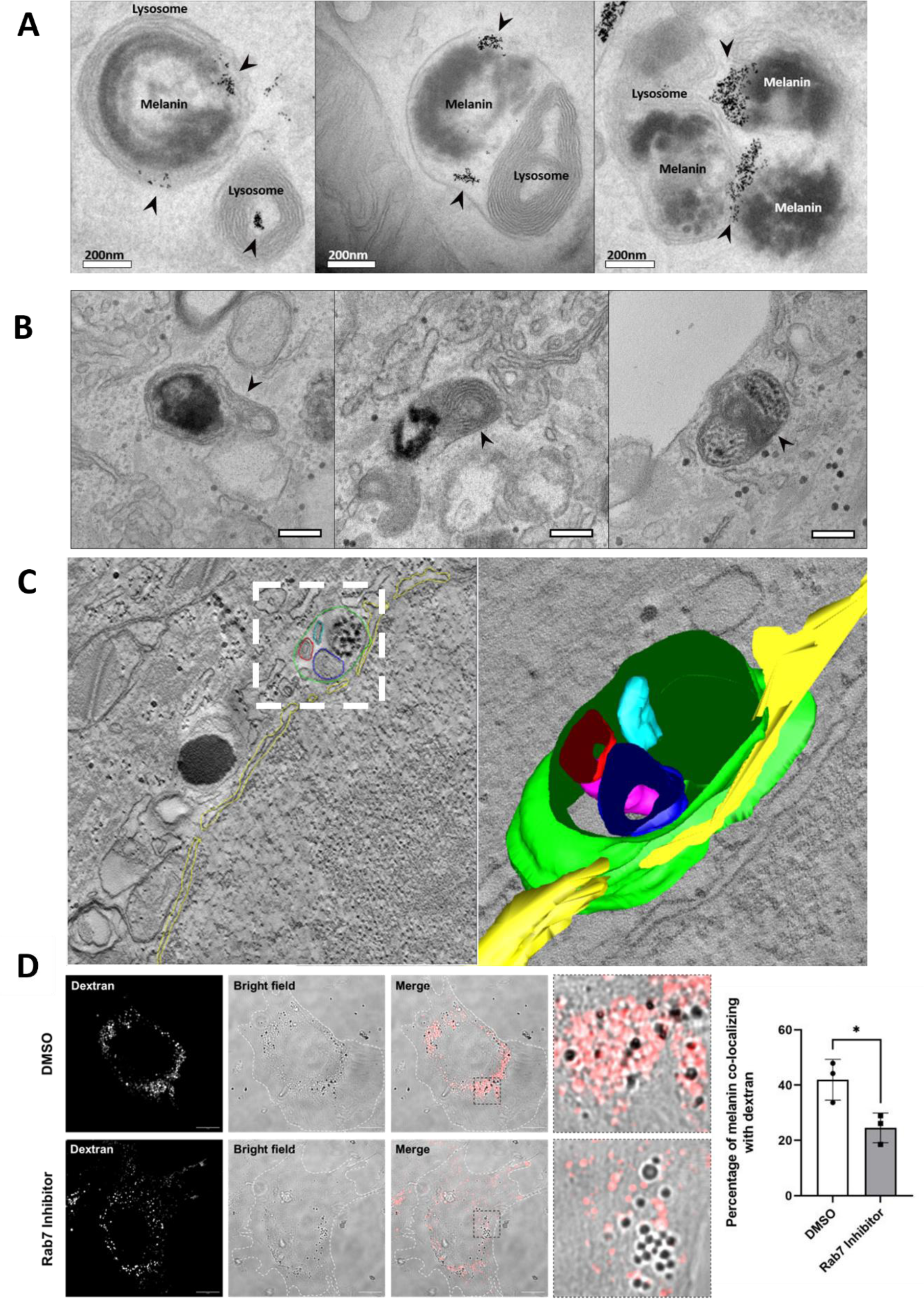
Melanin-containing phagosomes fuse with lysosomes, and the resulting melanokerasomes (MKSs) exhibit lysosome-like morphology. **(A)** XB2 keratinocytes were pulsed with BSA-gold conjugate (5 nm Au, arrows) for 2 hours and chased for 1 hour to load the lysosomal compartment, then pulsed with melanocores for 2 hours and fixed immediately after the pulse. Melanin-containing phagosomes can be observed fusing with gold-containing lysosomes when observed by TEM. Scale bars = 200 nm. **(B)** Examples of MKSs with lysosome-like morphology (arrows) are shown in reconstructed human pigmented epidermis keratinocytes. Scale bars = 200 nm. **(C)** A MKS positioned close to the nuclear envelope (yellow) was imaged 3-dimensionally by transmission electron tomography, revealing the presence of lysosomal lamellae (blue) and intralumenal vesicles (red, turquoise) enclosed within the same single membraned compartment (green) as the melanin. **(D)** XB2 keratinocytes were incubated with fluorescent dextran to load lysosomes, before incubating with a Rab7 inhibitor (CID1067700) and then feeding with melanocores. Co-localization between dextran and melanin was quantified. Scale bars = 10 μm. Quantification represents a minimum of 50 cells per condition.

It was previously reported that the recruitment of Rab-interacting lysosomal protein (RILP) by Rab7 and its consequent binding to the dynein-dynactin motor complex, promotes the emission of tubular protrusions that allow the fusion of late phagosomes and lysosomes (Harrison et al. 2003). This led us to inquire whether Rab7 could also mediate the fusion between melanin-containing phagosomes and lysosomes. To address this question, we performed a pulse-chase assay to load lysosomes with fluorescent dextran. Keratinocytes were simultaneously incubated with melanocores and treated with the Rab7 competitive inhibitor CID1067700 (Agola et al., 2012) or vehicle, and the co-localization between dextran and melanocores assessed by live-imaging. The results showed a significant decrease (∼41%) in the co-localization between melanocores and dextran in cells treated with the Rab7 inhibitor, indicating that Rab7 regulates the fusion between melanin-containing phagosomes and lysosomes (Figure 2D). Overall these results suggest that, after internalization by keratinocytes, melanin-containing phagosomes undergo maturation along the endocytic pathway, which culminates in their fusion with lysosomes, originating a specialized lysosomal compartment – the MKS.

### Melanokerasome juxtanuclear positioning is dependent on Rab7 and RILP

In keratinocytes, melanin accumulates in the juxtanuclear region. Since MKSs are specialized phagolysosomes, we hypothesized that the molecular machinery involved in the transport of MKSs to the juxtanuclear region of keratinocytes is shared with lysosomes. Lysosome positioning depends on their long-range transport along microtubules, which is regulated by the relative activity of the motor proteins dynein and kinesins (Matteoni & Kreis, 1987; Hollenbeck & Swanson, 1990; Harada et al, 1998; Pu et al, 2016). The anterograde transport of lysosomes is predominantly regulated by the interaction between Arl8b and the BLOC-1-related complex (BORC) (Pu et al., 2015). Arl8b-BORC complex can mediate the direct binding of lysosomes to kinesin-3 or the indirect binding of lysosomes to kinesin-1, through the recruitment of the Arl8b effector, Pleckstrin homology domain-containing family M member 2 (PLEKHM2, also known as SKIP) (Rosa-Ferreira & Munro, 2011; Khatter et al., 2015; Guardia et al., 2016). Accordingly, we observed that the overexpression of Arl8b promotes an accumulation of both LAMP-1-positive vesicles and MKSs at the periphery of keratinocytes, which suggests that the transport of MKSs in keratinocytes is regulated by the same molecular mechanisms as lysosomal positioning (Supplementary Figure S3).

On the other hand, the retrograde transport of lysosomes is mainly mediated by Rab7. After activation, Rab7 recruits its effector RILP, which binds to the light intermediate chain of dynein and to the p150^Glued^ subunit of dynactin, promoting the transport of lysosomes toward the juxtanuclear region of the cells (Jordens et al., 2001; Cantalupo et al., 2001). To assess whether Rab7 regulates the transport of MKSs in keratinocytes, we silenced either Rab7a and/or Rab7b and observed an impairment in the juxtanuclear clustering of MKSs in the cells silenced for Rab7a/b, when compared to control cells (Figure 3A and C). In addition, we found that the silencing of Rab7a/b promotes a dispersion of LAMP-1-positive vesicles. These phenotypes were also observed when we used the Rab7 inhibitor CID1067700 (Supplementary Figure S3). Since RILP promotes the interaction between Rab7 and the dynein/dynactin motor complex (Jordens et al., 2001; Cantalupo et al., 2001), we next assessed if RILP is involved in the transport of MKSs to the juxtanuclear region of keratinocytes. As expected, we observed a dispersion of MKSs, as well as LAMP-1-positive vesicles, in cells depleted of RILP (Figure 3B and D). These results suggest that Rab7 and its effector RILP mediate the retrograde transport of MKSs, to the juxtanuclear region of keratinocytes.

**Figure 3:**
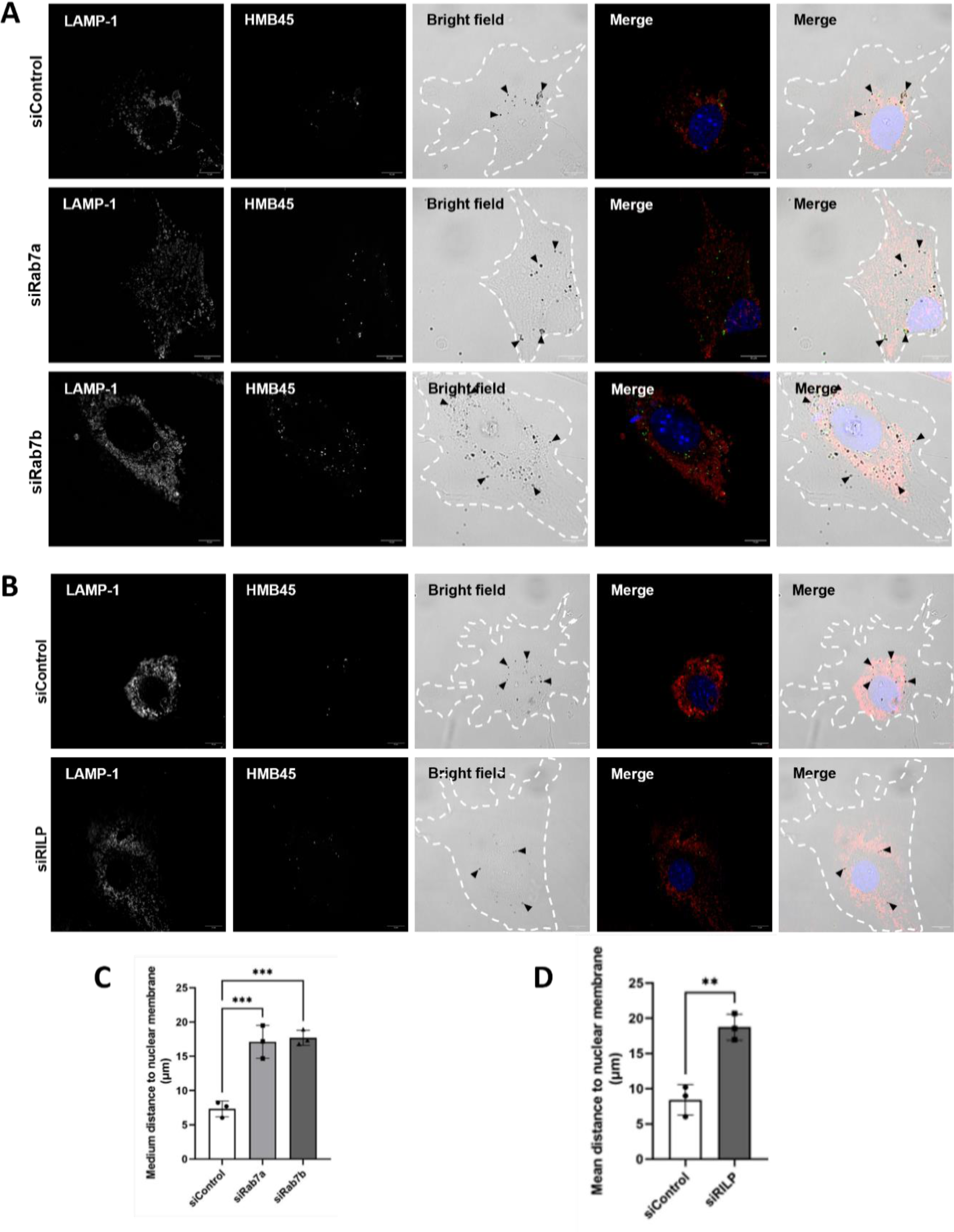
Impaired MKS juxtanuclear positioning upon modulation of lysosomal positioning machinery components. XB2 keratinocytes were transfected with non-targeting siRNA (siControl), siRNA targeting Rab7a (siRab7a), Rab7b (siRab7b) **(A)** or RILP (siRILP) **(B)**. Cells were then incubated with 0.1 g/L of melanocores for 4 hours and melanin was chased for another 20 hours. After the chase, cells were fixed and the distribution of melanocores inside keratinocytes was assessed by immunofluorescence, with the LAMP-1 (red) and HMB45 (green) antibodies. Nuclei were visualized by DAPI staining (blue). Scale bars = 10 μm. Arrowheads indicate the melanocores inside keratinocytes. Quantification of the mean distance of melanocores to the nuclear envelope after the silencing of Rab7a and Rab7b **(C)** or RILP **(D)**. p values (one-way ANOVA) were considered statistically significant when <0.001 (***) or when <0.01 (**). Plots show mean ± SD of three independent experiments.

### Melanin persists in a storage lysosome

Previous work from our group established that melanin persists in keratinocytes for at least 7 days (Correia et al., 2018). Therefore, we aimed to understand how melanin is able to evade degradation, despite fusion of melanin-containing phagosomes with lysosomes. Although previous studies appeared to point towards the arrested maturation hypothesis (Correia et al., 2018; Hurbain et al., 2018), a more recent unveiling of the lysosome complex dynamics (Bright et al., 1997; Bright et al. 2016; Barral et al., 2022) raised the possibility that MKSs exist in a hydrolase-inactive stage of the lysosome cycle after acquisition of melanin. Therefore, we measured lysosomal hydrolase activity by flow cytometry at different timepoints after melanocore internalization, using Magic Red Cathepsin B (MRCatB). MRCatB is membrane-permeable probe that fluoresces upon cleavage by Cathepsin B (CatB), and thus can be utilized as a specific readout of lysosome activity. We observed no significant differences in MRCatB fluorescence at 1h and 4h, when compared with control cells that were not fed melanin (Figure 4A). Conversely, at 24h after melanin internalization, we observed a significant decrease in the fluorescence of MRCatB, which suggests a reduction in lysosomal hydrolase activity caused by the presence of melanin inside lysosomes.

**Figure 4:**
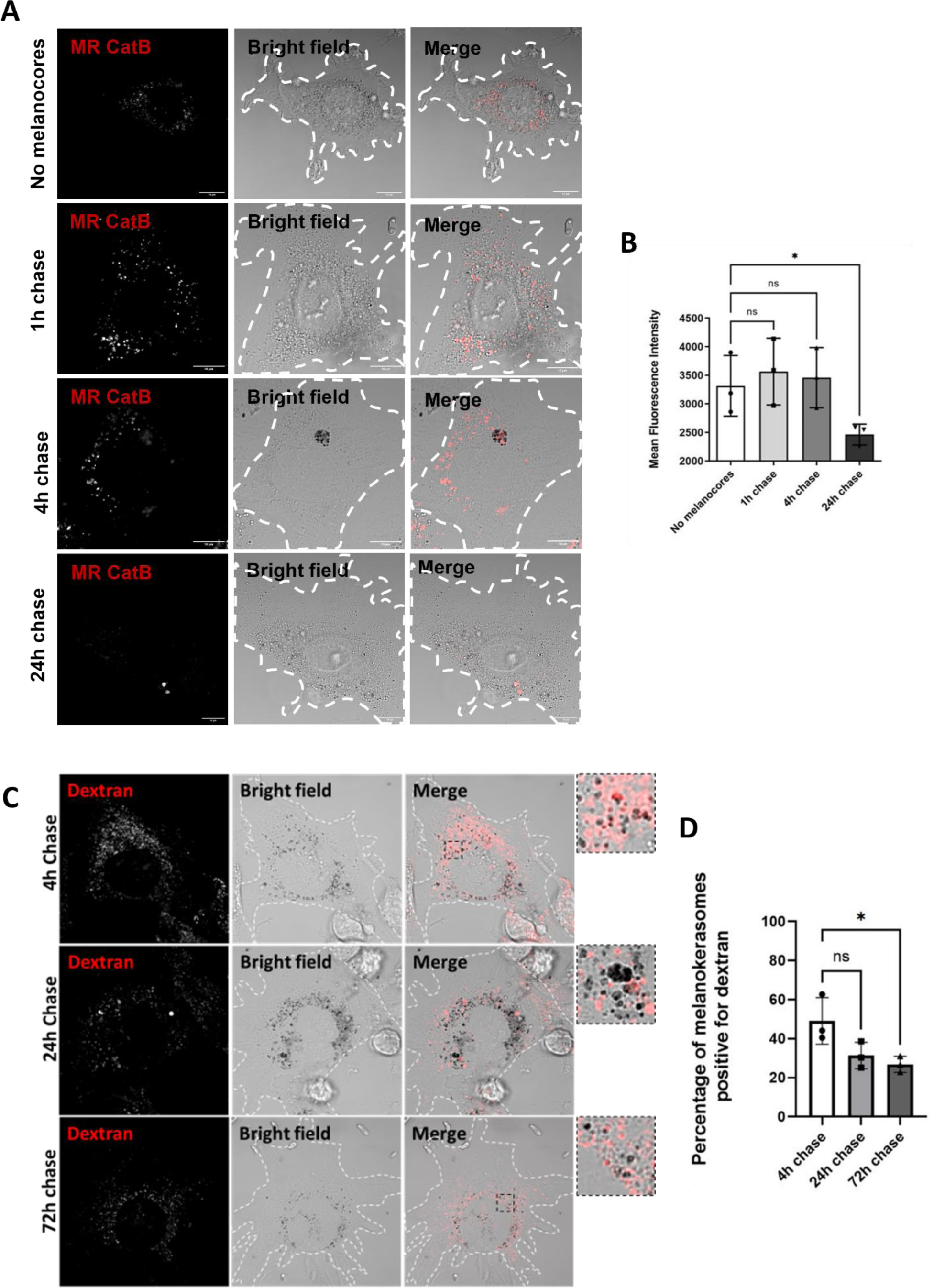
Melanokerasome (MKS) maturation is accompanied by a reduction in lysosomal hydrolase activity and decreased accessibility to fluid phase cargo. **(A and B)** XB2 keratinocytes were fed with melanocores, chased for varying periods of time (1h, 4h or 24h), and Cathepsin B activity assessed using Magic Red Cathepsin B probe (MR CatB). Representative confocal images of MR CatB fluorescence are shown in **(A)**, and quantitative analysis of CatB activity performed by flow cytometry is shown in **(B)**. Scale bars = 10 µm. p values (one-way ANOVA) were considered statistically significant when <0.05 (*) or non-significant (ns) when ≥0.05. Plots show mean ± SD of three independent experiments. **(C and D)** XB2 keratinocytes were fed with melanocores, chased for varying periods of time (4h, 24h or 72h), then pulsed and chased with dextran Alexa Fluor 568 to load the terminal endolysosomal compartment. The percentage of MKSs accessible to the newly-endocytosed dextran was measured through the co-localization between melanin (brightfield) and dextran. Scale bars = 10 µm. p values (one-way ANOVA) were considered statistically significant when <0.05 or non-significant (ns) when ≥0.05. Plots show mean ± SD of three independent experiments.

It has been suggested that lysosomes in certain stages of the lysosome cycle, or dysfunctional lysosomes, can be less fusogenic and thus less accessible to newly internalized material (Escrevente et al., 2021; Cardoso, Hall & Burgoyne et al., 2023). Therefore, we analyzed whether pre-formed MKSs are accessible to newly-internalized fluid-phase cargo. To do so, XB2 cells were fed with melanocores, chased for either 4h 24h or 72h, and then pulse-chased with fluorescent dextran. We observed no significant differences in MKS accessibility to dextran between 4h and 24h (Figure 4B). At 72h, we were able to detect a reduction in fluorescent punctae co-localizing with MKSs, which indicates that these mature compartments lose their ability to fuse with late endosomes/lysosomes (Figure 4B). Altogether, the observed decreases in lysosomal hydrolase activity and accessibility to the endolysosomal pathway at later timepoints suggest that these compartments drop out of the lysosome cycle as they mature, and therefore represent lysosomes in the terminal stage of their life cycle.

## Discussion

Herein, we unravel the cellular mechanisms underlying melanin processing and fate after transfer to keratinocytes to achieve efficient photoprotection. The results presented suggest that melanin-containing phagosomes mature along the endolysosomal pathway, fuse with lysosomes, and that melanin resides within a terminal lysosomal compartment, likely tethered to the nuclear membrane and inaccessible by the endolysosomal pathway (Figure 5). We observed a time-dependent and significant increase in the number of melanin-containing phagosomes associating with the late endosomal/lysosomal markers CD63 and LAMP-1 after melanocore internalization, in addition to presenting evidence that these compartments fuse with pre-existing lysosomes. This suggests that, after phagocytosis by keratinocytes, melanin-containing phagosomes undergo maturation along the canonical endocytic pathway, which is contrary to the alternative hypothesis that the MKS represents a maturation-arrested compartment. Notably, our results showed only a small decrease in the number of EEA-1-positive vesicles surrounding melanin-containing phagosomes with time. Importantly, recent studies found that EEA-1 only partially co-localizes with Rab5-positive vesicles and is also recruited to late endosomes (van der Beek et al., 2022), suggesting a more widespread distribution of this marker than previously accepted. Since MKSs originate from the fusion between melanin-containing phagosomes and lysosomes, this led us to hypothesize that transport of the former to the juxtanuclear region of keratinocytes relies on similar mechanisms and machinery that regulate lysosome positioning. In fact, we observed that the overexpression of Arl8b induces the dispersion of melanin throughout the cytoplasm of keratinocytes, which is in agreement with this protein regulating the kinesin-dependent trafficking of lysosomes toward the periphery of the cells (Rosa-Ferreira & Munro, 2011; Niwa et al., 2016).

**Figure 5:**
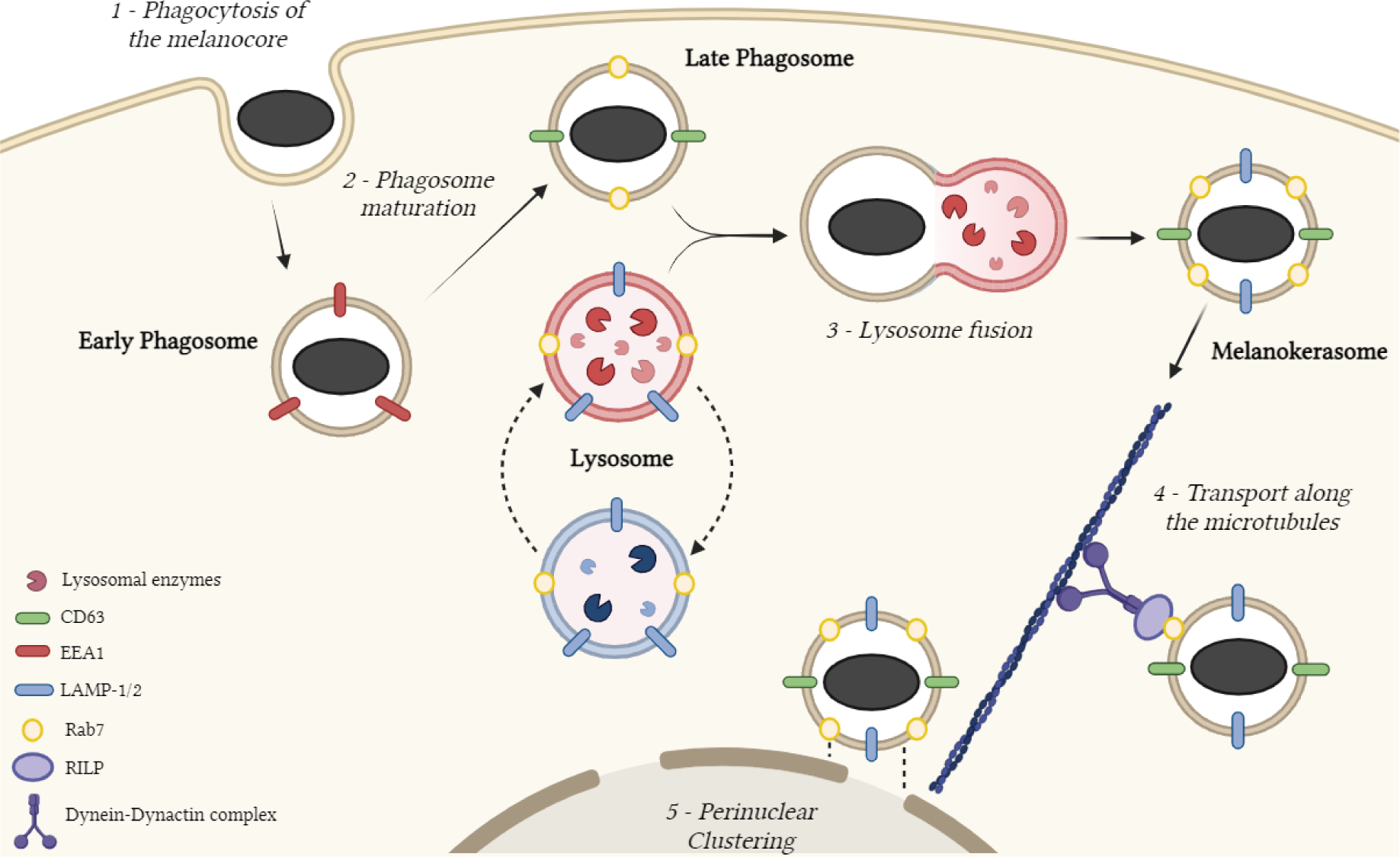
Schematic representation of the working model of MKS formation and maturation. Melanocores secreted by melanocytes are phagocytosed by keratinocytes **(1)**, and sequentially acquire early and late endolysosomal markers **(2)** before fusion with lysosomes to form melanokerasomes **(3)**. These are transported towards the nucleus using canonical lysosomal positioning machinery **(4)** and cluster around the apical pole of the nucleus to form photoprotective parasol-like supranuclear caps **(5)**.

We also hypothesized that MKSs utilize lysosomal trafficking machinery as an efficient redundancy mechanism to form the terminal compartment and establish juxtanuclear positioning, which could play a role the formation of supranuclear caps. Indeed, we found that the silencing of Rab7 or RILP significantly impairs the retrograde transport of MKSs inside keratinocytes. Strikingly, this phenotype was observed not only with the silencing of Rab7a, but also Rab7b. While Rab7a plays a well-known role in endosome maturation and lysosomal trafficking, Rab7b was described to control the late endosome-to-TGN transport (Progida et al.., 2010). Nevertheless, a recent study reported that melanosomes internalized by keratinocytes are stored in compartments that are positive for Rab7b (Marubashi & Fukuda, 2020).

We also provide evidence that melanin-loaded lysosomes of keratinocytes have reduced lysosomal hydrolase activity after prolonged chase periods, and that MKSs become less accessible to newly-endocytosed fluid phase cargo. Considering that melanin is known to be a dense polymer that is resistant to degradation by biochemical means (Borovanský & Elleder, 2003), and to our knowledge no concrete evidence demonstrating melanin degradation in mammalian cells has been put forward, we propose that MKSs represent a storage lysosome. Indeed, it would be energetically favorable for cells to maintain MKSs in this state and ensure that melanin persists for long periods without significant degradation. The characteristics of the MKS we found, namely exhibiting lysosome markers, eluding connections with the endo/phagocytic pathway, being weakly acidic/degradative and retaining undigested cargo (melanin), have been ascribed to dysfunctional lysosomes in pathological processes, from hereditary lysosomal storage diseases to neurodegenerative diseases (Nixon, 2020; Burrinha et al., 2023). This is also the case in the formation of lipofuscin-like auto-fluorescent granules in retinal pigment epithelial cells implicated in age-related macular degeneration (Escrevente et al., 2021). Therefore, we suggest that undigested cargo material, the loss of hydrolytic activity and increased pH, lead MKSs to drop-out of the lysosome cycle and become storage (dysfunctional) lysosomes. MKSs could thus represent the physiologically beneficial parallel to lysosome dysfunction. Understanding further the mechanisms leading to the creation of storage lysosomes may help elucidate critical pathogenic mechanisms in age-related diseases and uncover novel therapeutic approaches targeting the lysosome and endolysosomal trafficking machinery.

## Materials and Methods

### Cell culture and Reconstructed Human Pigmented Epidermis

XB2 murine keratinocytes were cultured in Dulbecco’s Modified Eagle Medium (DMEM) (Gibco/Invitrogen), supplemented with 10% fetal bovine serum (FBS), 2 mM L-Glutamine, 100 U/mL penicillin and 100 U/mL streptomycin. MNT1 human melanoma cells were cultured in DMEM, supplemented with 10% FBS, 2 mM L-Glutamine, 1% Non-Essential Amino Acids (NEAA) (Gibco/Invitrogen), 100 U/mL penicillin and 100 U/mL streptomycin. Cells were maintained at 37°C, 10% CO_2_ and 95% humidity. Reconstructed Human Pigmented Epidermis (RHPE) were formed as previously described (Zoio et al., 2021; Hall and Lopes-Ventura et al., 2022). Briefly, melanocytes and keratinocytes were seeded upon a polycarbonate membrane, and keratinocyte differentiation and stratification induced by establishing an air-liquid interface.

### Isolation of melanocores

MNT1 melanocytes were cultured for 7 days, in 150 cm^2^ T flasks (Corning), in complete growth medium. Afterwards, the culture medium, referred from now on as conditioned medium, was collected and concentrated by sequential centrifugations of 30 minutes at 3,800 x *g*, in a Vivaspin Centricon (Sigma), with a pore size of 300 kDa. The absorbance value of the melanocore preparation was measured using a Nanodrop 2000 (Thermo Scientific) at 340 nm and the concentration of melanin was calculated using a calibration curve developed in-house: [Absorbance = 1.8546 x Concentration (g/L) - 0.0422].

### siRNA silencing

XB2 keratinocytes were seeded in 24-well plates, at a confluency of 4×10^4^ cells/well. Twenty-four hours later, 50 nM of siRNA pools (Thermo Scientific, Supplementary Table 1) were added to 50 μL of Opti-MEM (Gibco/Invitrogen). Simultaneously, 1.5 μL of Dharmafect 4 (Dharmacon) was diluted in 50 μL of Opti-MEM. Both mixtures were incubated at room temperature (RT) for 5 minutes and then combined, mixed gently, and incubated for 20 minutes at RT. Culture medium was removed before transfection and 100 μL of siRNA mixture was added to 400 μL of Opti-MEM. Cells were then incubated in normal culture conditions and, 24 hours later, the transfection medium was replaced with complete growth medium. A non-targeting siRNA pool (Thermo Scientific, Supplementary Table 1) was used as a control. Efficient silencing was validated by quantitative real-time PCR (qRT-PCR) analysis (Supplementary Figure S1).

### Plasmid transfection

XB2 cells were seeded in 24-well plates, at a confluency of 4×10^4^ cells/well. Twenty-four hours after seeding, a mixture of 1 μg of plasmid (Supplementary Table 2) in 100 μL of Opti-MEM was combined with 1.5 μL of TurboFect (Thermo Scientific), diluted in 100 μL of Opti-MEM, according to the manufacturer’s instructions. Cells were then incubated for 24 hours, in normal culture conditions, and the medium was changed to 500 μL of complete growth medium. The plasmids used are described in the Table below.

### Real-time quantitative PCR

Total RNA was isolated using the RNeasy Mini kit (Qiagen), according to the manufacturer’s instructions. Superscript® II DNA synthesis kit (Invitrogen) was used to reverse-transcribe 500 ng to 1 µg of total RNA and synthesise cDNA. qRT-PCR reactions were performed using a Roche LightCycler and the Roche SybrGreen Master Mix reagent. The reaction mixture was composed of 5 μL of SybrGreen, 4 μL of cDNA and 10 μM of specific primers (Supplementary Table 3), for a total reaction volume of 10 μL. For each protein, gene expression was calculated relative to control wells, with the LightCycler96 software (Roche). Gapdh was used as a housekeeping gene. Quadruplicates were used for each gene.

### Immunofluorescence

Cells were fixed with 4% PFA in 1X PBS for 20 minutes. The excess of fixative was removed by washing with 1X PBS for 10 minutes and permeabilization and blocking were performed with a solution of 1% (m/v) BSA (Sigma) and 0.05% (m/v) saponin (Sigma) in 1X PBS, for 30 minutes. The cells were then incubated with diluted primary antibodies (Supplementary Table 4) for 1 hour and 30 minutes, washed 3 times with 1X PBS and subsequently incubated for 1 hour with Alexa fluorophore-conjugated secondary antibodies, diluted 1:500 in blocking buffer (Supplementary Table 4). Unbound antibodies were removed with extensive washing with 1X PBS. In the specific case of the immunostaining with HMB45 and anti-LAMP-1 antibodies, the latter was added after the incubation with the secondary antibody, and after extensive washing with 1X PBS, to avoid any unspecific cross-reactions. Coverslips were then mounted on glass slides with the Fuoromount-G^TM^ mounting media containing DAPI (Invitrogen), to stain the nuclei of the cells. All antibody incubations and washes were performed with a solution of 1% (m/v) BSA and 0.05% (m/v) saponin and 1X PBS, respectively. All steps were performed at RT.

### Cathepsin B activity assay

Cathepsin B (CatB) activity was detected and measured using Magic Red CatB (MR CatB) substrate from ImmunoChemistry Technologies, which fluoresces upon cleavage by active CatB. Briefly, MR CatB was added to cells for 30 minutes prior to imaging/fluorescence measurement at a concentration of 673 nM, in normal culture conditions, and fluorescence was measured by flow cytometry. For each condition, a parallel glass coverslip of cells with identical treatment was imaged by live cell confocal microscopy to provide representative images and to confirm probe specificity.

### Flow cytometry

XB2 cells incubated with Magic Red were washed with PBS and flow cytometry (FC) buffer [1% FBS (v/v) and 2 mM EDTA in PBS], centrifuged at 300 x g for 5 minutes and resuspended in FC buffer. Data acquisition was performed in a FACS CANTO II flow cytometer (BDBiosiences). At least 30,000 cells were acquired per condition using BD FACSDivaTM software (Version 6.1.3, BD Biosciences). Data analysis was performed in FlowJo (Version 10, BD Biosciences).

### Lysosome loading with dextran

XB2 keratinocytes were plated on Nunc Lab-Teks (Thermofisher), at a confluency of 1.5×10^4^ cells/well. The following day, 1 mg/mL of dextran Alexa Fluor 568 (10 kDa; Invitrogen) was fed to the cells for 4 hours and chased for 20 hours, in order to accumulate in lysosomes. Then, keratinocytes were incubated with 0.1 g/L of melanocores for 4 hours, chased for 20 hours and the co-localization between dextran and melanin was assessed by live-cell imaging. To determine if MKSs are accessible to fluid phase cargo, XB2 cells were pulsed with melanocores for 1 hour on ice, then chased for 4, 24 or 72 hours. While the cells were being chased, they were also pulsed with dextran for 2 hours and chased for another 2 hours. After this, live imaging of the cells was performed by confocal microscopy.

### Preparation and loading of BSA-gold conjugate

After performing a stabilization test to determine the amount of protecting bovine serum albumin (BSA, Sigma) required to stabilize a fixed volume of 5 nm gold colloid suspension (BBI Solutions), the appropriate volumes of BSA in 2 mM borax and 20 ml gold colloid suspension were combined. After gentle agitation for 5 minutes, the preparation was secondarily stabilized with a solution of 10% BSA in 2 mM borax to a final concentration of 1% BSA. The preparation was centrifuged at 45,000 rpm (205,250 x g) for 25 minutes and the fluid pellet containing the BSA-gold conjugate was extracted. The final fluid pellet formed from 22.5 ml of the starting gold colloid suspension was diluted 1:7 in media and fed to the cells, as with dextran.

### Melanin distribution quantification

XB2 keratinocytes transfected with siRNAs (Supplementary Table 1) or incubated with 25 nM of a Rab7 inhibitor (CID-1067700, Sigma-Aldrich) were then fed with 0.1 g/L of melanocores for 4 hours and melanin was chased for another 20 hours. After the chase, cells were fixed and the distribution of late endosomes/lysosomes and melanin-containing compartments inside keratinocytes was assessed by immunofluorescence, with anti-LAMP-1 and HMB45 antibodies (Supplementary Table 4), respectively. Brightfield microscopy was also used to identify dark melanin-containing compartments. Melanin distribution was assessed by an ImageJ macro developed and kindly shared by Silvia Benito-Martínez and Cédric Delevoye (Institut Curie). Nuclei were counted to ensure similar cell confluence in all samples.

### Fluorescence Microscopy

Confocal imaging was performed with a Zeiss LSM980 confocal microscope, equipped with a 40x 1.1 NA water objective and 63x 1.4 NA oil objective, and live imaging was performed at 37°C in HEPES buffer/media. Super-resolution images were acquired using the Airyscan 2 detector (Zeiss).

### Transmission electron microscopy

All reagents and materials were purchased from Electron Microscopy Sciences unless otherwise stated. Specimens were fixed in 2% paraformaldehyde, 2% glutaraldehyde in 0.1 M sodium phosphate buffer overnight at 4°C. Specimens were post-fixed for 1 hour on ice using 1% osmium tetroxide and 1.5% potassium ferrocyanide in distilled water, and then incubated with 1% tannic acid in distilled water for 30 minutes at RT, before dehydrating with a series of increasing ethanol concentrations (70%, 90%, 2x 100%) and infiltrating/embedding in Epon resin (EMbed 812). After polymerizing resin at 65⁰C overnight, glass was removed by immersion in liquid nitrogen, and cell monolayers were sectioned ‘*en face*’ at 70 nm thickness using a Reichart Ultracut S ultramicrotome (Leica) and a diamond knife (Diatome), and sections collected on copper mesh grids. Sections were post-stained with 2% uranyl acetate in 70% methanol followed by Reynold’s lead citrate, and imaged using an FEI Tecnai G2 Spirit BioTWIN TEM equipped with an Olympus-SIS Veleta CCD camera. Tilt series for electron tomography were acquired using Serial EM software, tomogram generation was performed by back-projection in IMOD, and tomogram segmentation and modelling was performed in 3dmod.

### Image and statistical analysis

All images for this manuscript were analyzed using ImageJ software. A minimum of three independent experiments were performed as replicates, with at least 50 cells imaged and quantified per condition/replicate. Graphic representation and statistical analysis were performed in Prism (GraphPad Software Inc.). Student’s t-test and one-way ANOVA were used to analyze the p value of differences, between 2 or more conditions, respectively.

## Acknowledgements

We thank Prof. Paul Luzio (Cambridge Institute for Medical Research) for the critical reading of the manuscript. We also thank Dr. Silvia Benito-Martinez and Dr. Cédric Delevoye for sharing the ImageJ macro prior to publication, and Dr. Jaime Mota (Faculdade de Ciências e Tecnologia, Universidade NOVA de Lisboa) for the kind gift of the Arl8b-GFP construct. We also thank the Microscopy, Cell Culture and Flow Cytometry facilities at NMS Research, and the Electron Microscopy Facility at Instituto Gulbenkian de Ciência. This work was supported by iNOVA4Health (UIDB/04462/2020 and UIDP/04462/2020) and by the Associated Laboratory LS4FUTURE (LA/P/0087/2020), two programs financially supported by *Fundação para a Ciência e Tecnologia* (FCT)/*Ministério da Ciência, Tecnologia e Ensino Superior*, as well by FCT grant PTDC/BIA-CEL/29765/2017. MVN and JC were supported by PhD fellowships from FCT (PD/BD/137442/2018 and PD/BD/136905/2018, respectively).

---

**Figure S1:**
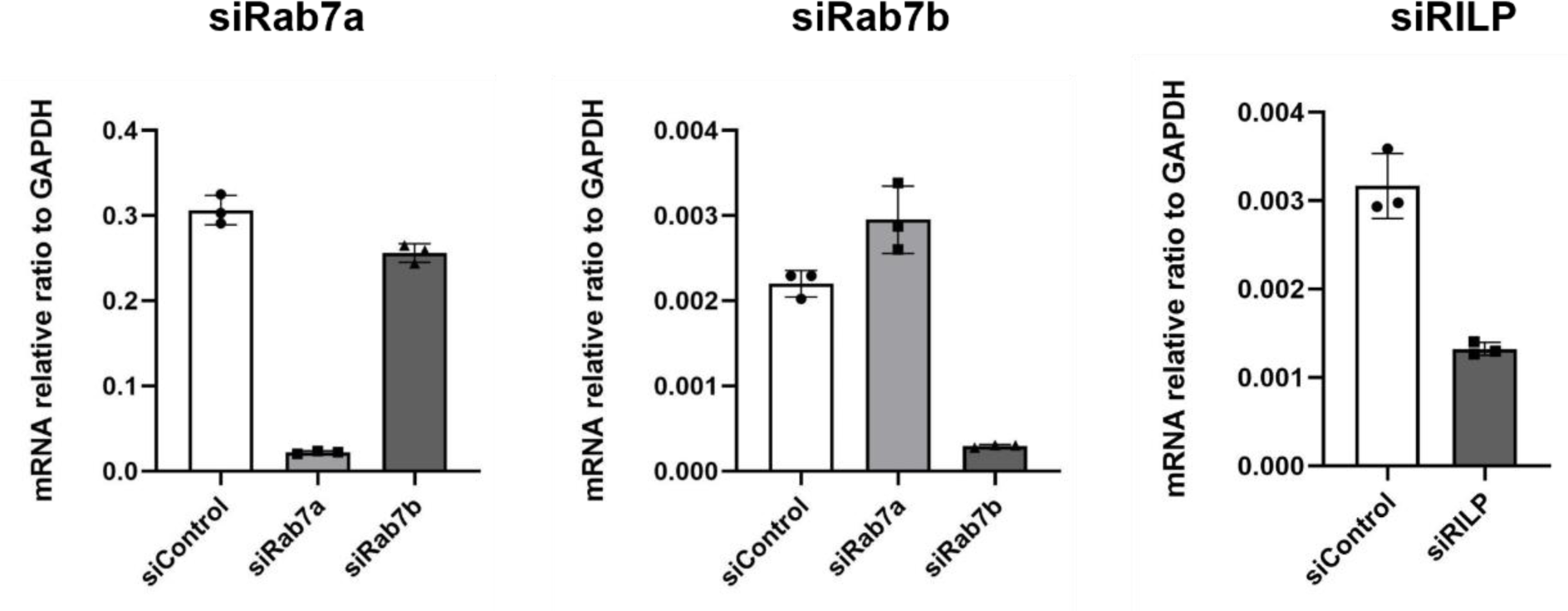
Efficiency of siRNA silencing, assessed by qRT-PCR measurement of mRNA levels. XB2 keratinocytes were transfected with non-targeting siRNA (siControl) and siRNA targeting Rab7a (siRab7a), Rab7b (siRab7b) or RILP (siRILP). The levels of silencing were measured by qRT-PCR, 72 hours after transfection.

**Figure S2:**
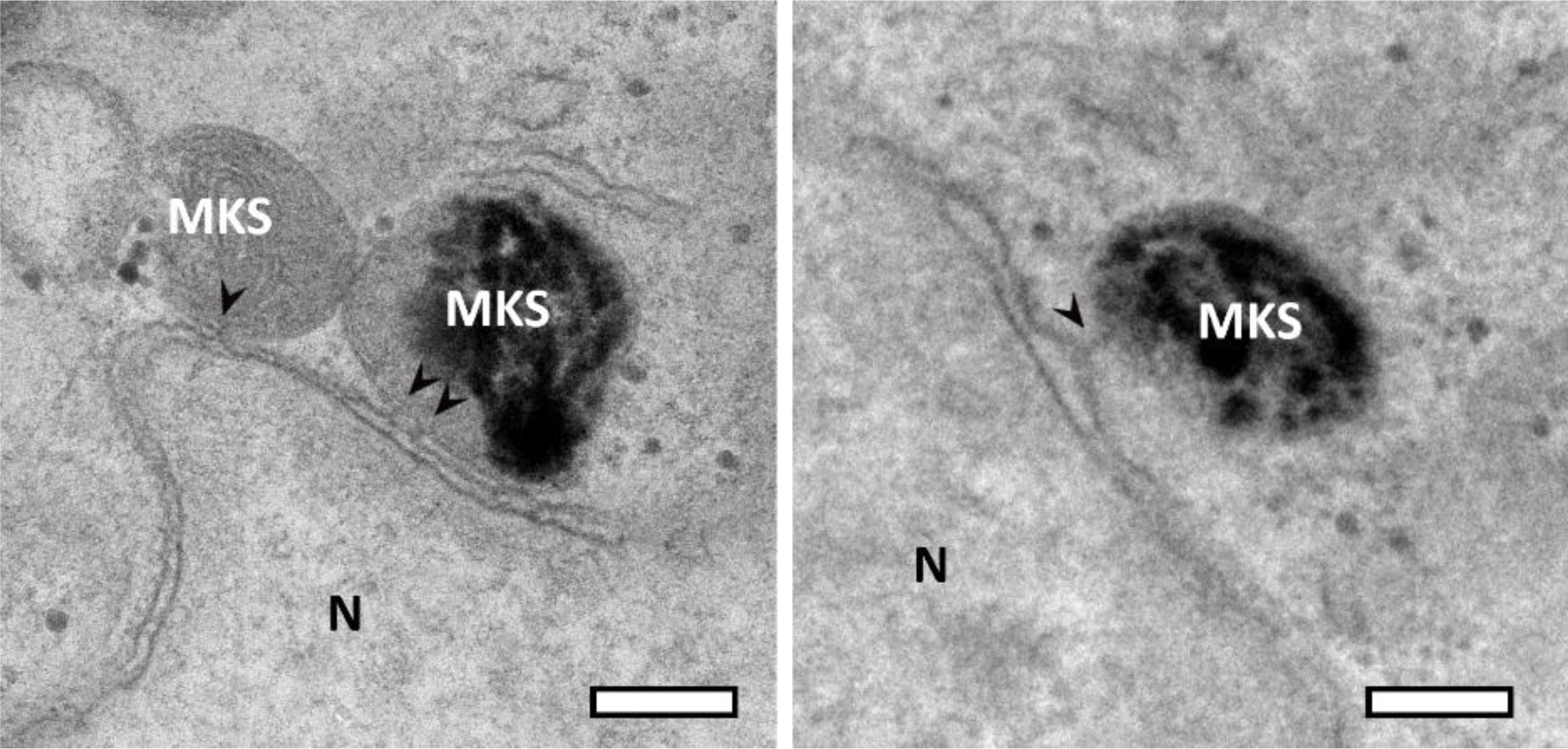
Melanokerasome-nucleus tethers. Melanokerasomes (MKS) in close apposition with the nucleus (N) within keratinocytes of Reconstructed Human Pigmented Epidermis were observed by TEM. Areas of electron density that could represent protein tethers between the outer nuclear membrane and the MKS limiting membrane are indicated by arrowheads. Scale bars = 200 nm.

**Figure S3:**
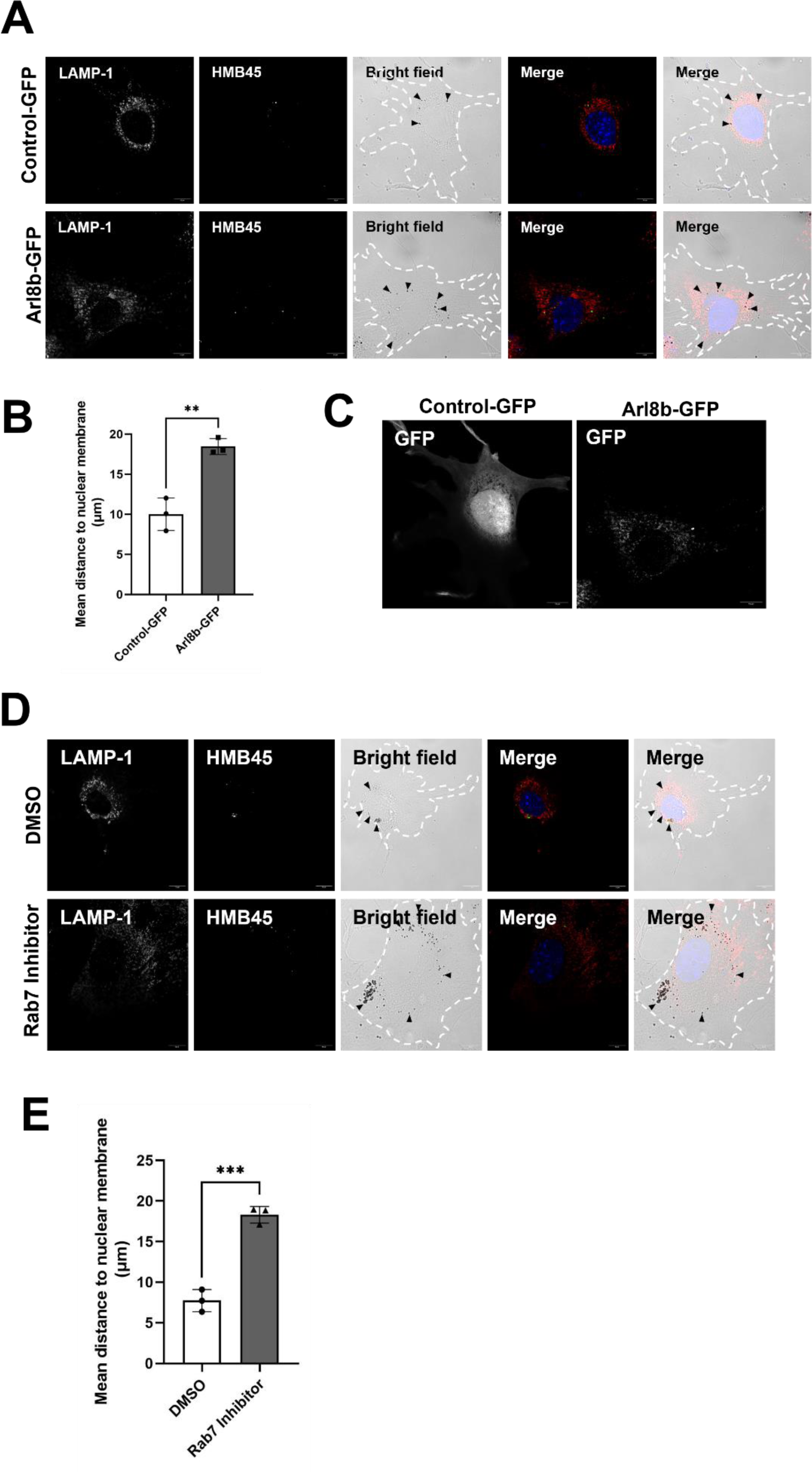
: Melanin juxtanuclear positioning is ablated upon Arl8 overexpression or Rab7 inhibition. **(A)** XB2 keratinocytes were transfected with non-targeting plasmid (Control-GFP) or plasmid to overexpress Arl8b (Arl8b-GFP). Cells were then incubated with 0.1 g/L of melanocores for 4 hours and melanin was chased for another 20 hours. After the chase, cells were fixed and the distribution of melanocores inside keratinocytes was assessed by immunofluorescence. LAMP-1 (red) and HMB45 (green) antibodies were used to label late endosomes/lysosomes and PMEL17, respectively. Nuclei were visualized by DAPI staining (blue). Arrowheads indicate the melanocores inside keratinocytes. **(B)** Quantification of the mean distance of melanocores to the nuclear envelope. **(C)** Transfection efficiency was assessed by GFP expression. Scale bars = 10 μm. **(D)** XB2 keratinocytes were treated with either the vehicle drug (DMSO) or with 25 μM of Rab7 inhibitor. Simultaneously, cells were incubated with 0.1 g/L of melanocores, for 4 hours, and melanin was chased for 20 hours. Afterwards, cells were fixed and the distribution of melanocores inside keratinocytes was assessed by immunofluorescence. LAMP-1 (red) and HMB45 (green) antibodies were used to label late endosomes/lysosomes and PMEL17, respectively. Nuclei were visualized by DAPI staining (blue). Scale bars = 10 μm. Arrowheads indicate the melanocores inside keratinocytes. **(E)** Quantification of the mean distance of melanocores to the nuclear envelope. p values (t-test) were considered statistically significant when <0.01 (**) or <0.001 (***) and non-significant (ns) when ≥0.05. Plots shows mean ± SD of three independent experiments.

**Table S1:** siRNAs used for silencing.

| siRNA | Species | Reference |
| --- | --- | --- |
| siControl | Mouse | D-001206-13 |
| siRab7a | Mouse | M-040859-01 |
| siRab7b | Mouse | M-053310-01 |
| siRilp | Mouse | M-063866-01-0005 |

**Table S2:** Plasmids used for overexpression.

| Overexpressed protein | Plasmid backbone | Source |
| --- | --- | --- |
| GFP | pENTR-GFPC1 | SicGen |
| Arl8b-GFP | pEGFP-N1 | Dr. Jaime Mota |

**Table S3:**
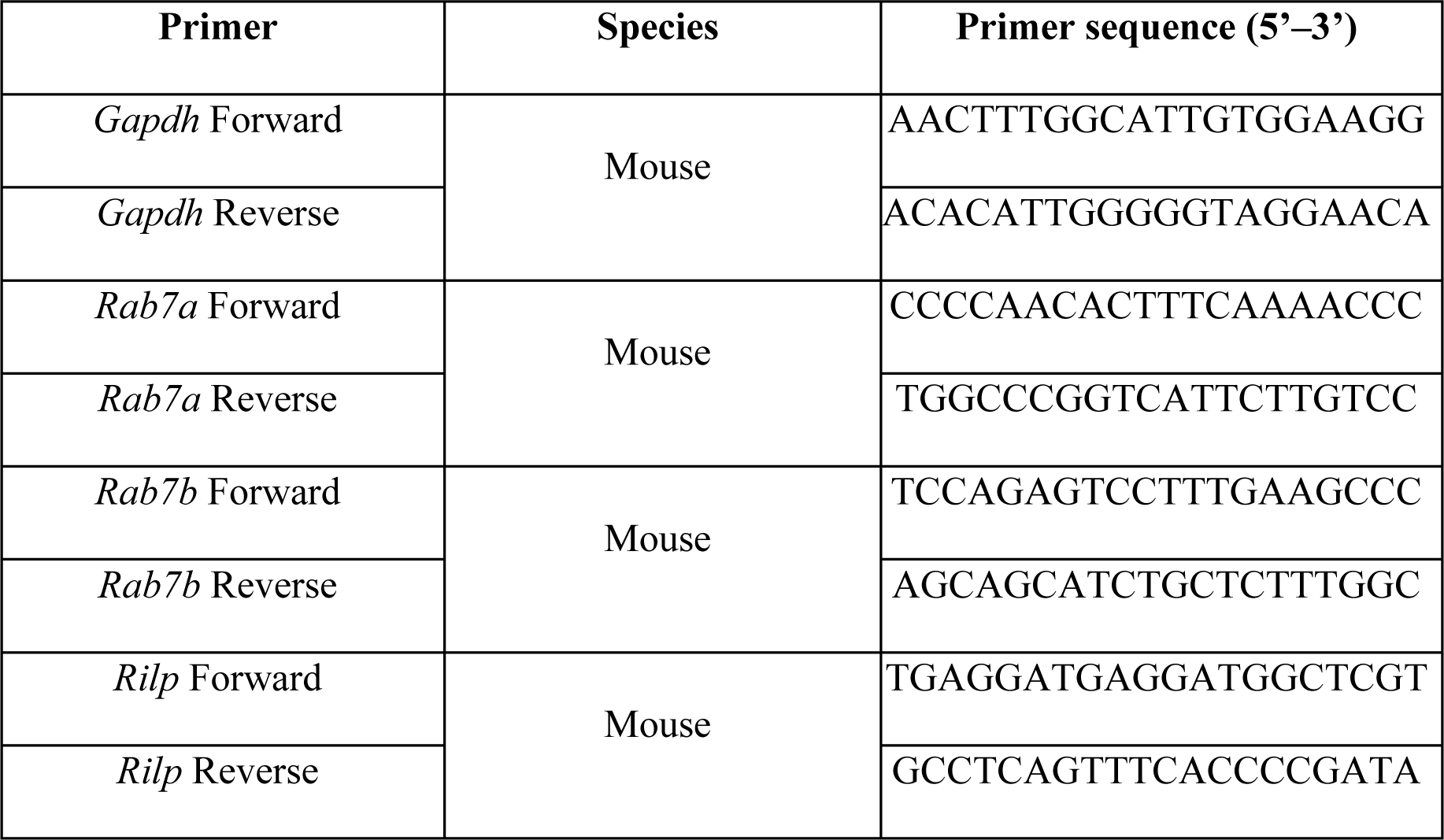
Primers used for qRT-PCR.

**Table S4:** List of primary and secondary antibodies.

| **Antibody** | **Host species** | **Brand/Ref.** | **Dilution** |
| --- | --- | --- | --- |
| Anti-CD63 | Mouse | MBL/R562 | 1:250 |
| Anti-EEA-1 | Mouse | BD Biosciences/610456 | 1:100 |
| HMB45 (anti-PMEL) | Mouse | Dako/M0634 | 1:250 |
| Alexa 647 anti-LAMP-1 | Mouse | Santa Cruz/ Sc19992 | 1:500 |
| Alexa Fluor 488 anti-mouse | Goat | Invitrogen/A11001 | 1:500 |
| Alexa Fluor 555 anti-mouse | Donkey | Invitrogen/A31570 | 1:500 |

